# FlexBench: *In Silico* and *In Vitro* Benchmarking of Structure-Based AI for Small Molecule Drug Design

**DOI:** 10.64898/2026.11.24.769037

**Authors:** Kyle Swanson, Dabrick Brill, Raktim Mitra, Swetha Srinivasan, Marion Lanier, Esther Martinborough, Sylvain Gariel, Alexander S. Powers

**Author notes:** 40th Conference on Neural Information Processing Systems (NeurIPS 2026). Workshop: AI for Drug Discovery: Bridging the Translation Gap.

## Abstract

Structure-based AI methods for small molecule drug discovery (SMDD) use AI to design molecules that bind to a protein given its 3D structure, with the goal of accelerating the discovery of new small molecule therapeutics. However, these methods have not been carefully benchmarked against each other or traditional methods for SMDD. To fill this gap, we present FlexBench, a benchmark of recent structure-based AI methods for SMDD, including our internal model SAGE-Flex and BoltzMol-1, compared to docking-based methods on four internal GPCR targets. Unlike prior benchmarks, FlexBench holds the compute budget and chemical space constant, making differences in performance attributable to the methods rather than their resources. Furthermore, we evaluated designs not just *in silico* but also *in vitro*, with 276 compounds synthesized and tested, which is one to two orders of magnitude more than typical prospective studies. While all methods had tradeoffs across the *in silico* metrics, the structure-based AI methods designed molecules in regions of chemical space that were unexplored by docking-based methods, thereby unlocking new molecular scaffolds. Of these methods, SAGE-Flex best combined that unique chemistry with a strong *in silico* profile, resulting in a hit rate of 12% *in vitro* versus 3% for BoltzMol-1. FlexBench thus effectively evaluates the strengths and weaknesses of structure-based AI methods for SMDD.

## 1 Introduction

Small molecule drug discovery (SMDD) has relied on established computational approaches such as virtual screening via molecular docking [1–3] to identify compounds that bind to a target protein. These methods can search increasingly large chemical libraries [4], especially when accelerated by artificial intelligence (AI) [5, 6]. More recently, advances in AI have produced a new class of structure-based AI models that use the 3D structure of proteins to design molecular binders without relying on docking. These structure-based AI models design molecules in various ways, from co-folding models that evaluate pre-specified molecules by predicting and scoring the 3D structure of a protein-ligand complex [7] to generative models that design molecules *de novo* conditioned on a known protein structure [8, 9]. Outside of SMDD, structure-based AI methods have achieved striking success in designing proteins and biologics, including antibodies, nanobodies, enzymes, and other protein binders [10–18]. However, structure-based AI methods have not yet been widely benchmarked for SMDD. To our knowledge, only a few benchmarks compare structure-based AI methods to docking-based methods *in silico*, and even fewer evaluate any molecules *in vitro*.

Here, we present FlexBench (Figure 1), a benchmark that evaluates structure-based AI methods for SMDD *in silico* and *in vitro*. FlexBench includes two structure-based AI methods: our internal generative model SAGE-Flex and BoltzMol-1 [7], an adaptation of the popular Boltz-2 structure prediction model [14] for SMDD. FlexBench compares these two methods against AI-accelerated docking and virtual screening on four G protein-coupled receptor (GPCR) targets from our internal programs. GPCRs are an important class of drug targets, with 36% of approved drugs targeting GPCRs across a multitude of disease areas [19], making them a good test case for structure-based AI.

**Figure 1:**
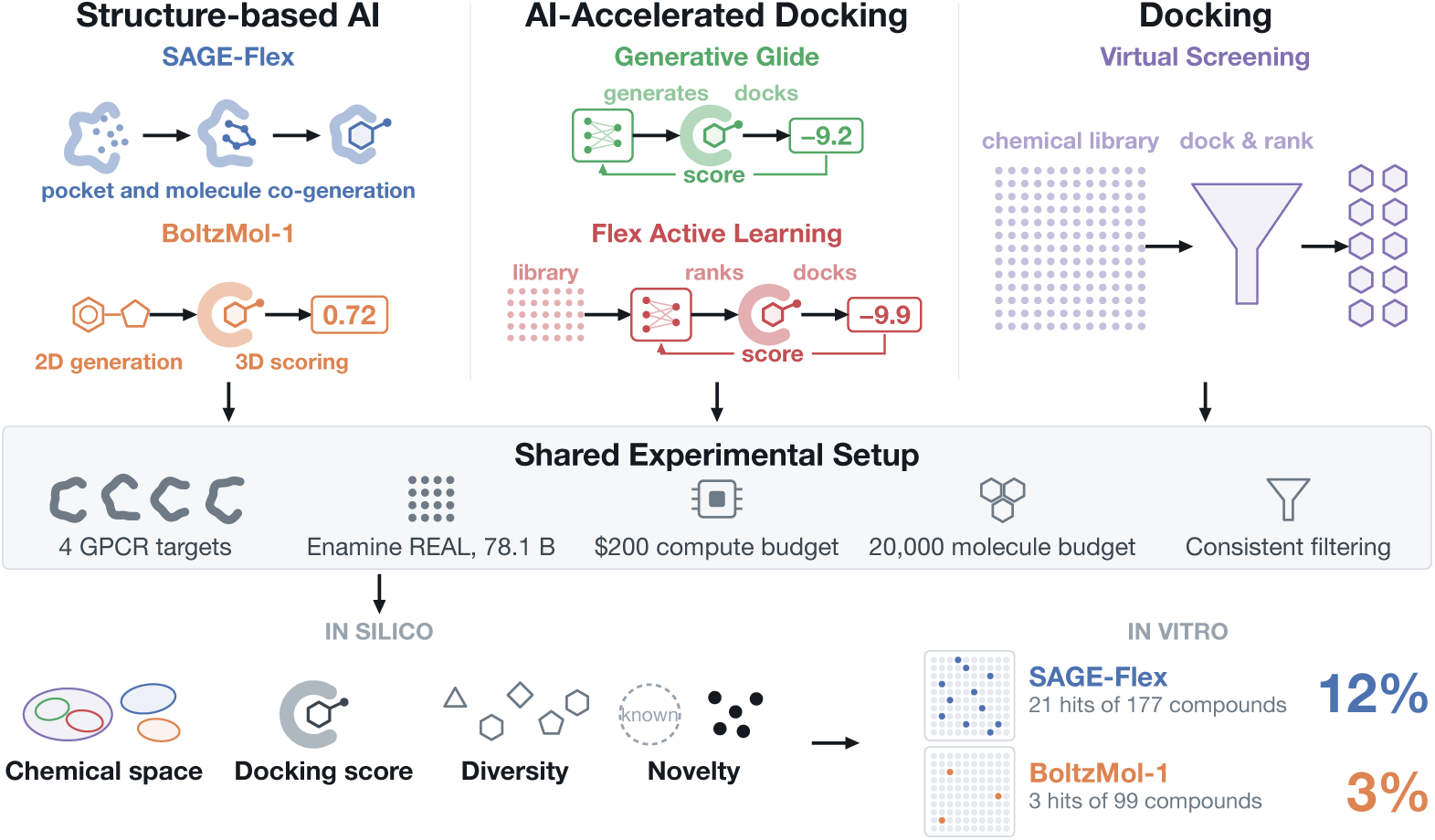
Overview of FlexBench. FlexBench compares structure-based AI methods for small molecule drug design to AI-accelerated docking and traditional docking (virtual screening) using a shared experimental setup with both *in silico* and *in vitro* results.

Previous benchmarks of AI methods for SMDD are generally limited to academic rather than industrial methods, use inconsistent compute budgets and an unrestricted chemical space, and score designs only on imperfect computational proxies rather than validating them *in vitro* [20–27]. FlexBench addresses these limitations by (1) including recent methods from both academia and industry, (2) fixing the compute budget [28, 29] and the chemical space to isolate differences to the methods themselves, and (3) prospectively testing top designs from structure-based AI methods *in vitro* at scale, with 276 compounds synthesized and tested across three targets. This is one to two orders of magnitude larger than typical prospective studies [30–32] and uniquely enables a head-to-head comparison of hit rates across structure-based AI models on the same targets under the same experimental conditions.

FlexBench showed that across all four targets, structure-based AI methods designed molecules in regions of chemical space that were distinct from docking-based methods, thereby unlocking new molecular scaffolds. However, no method dominated on all metrics, with each making a distinct tradeoff between its region of chemical space, docking scores, and diversity. Among the structure-based AI methods, SAGE-Flex outperformed BoltzMol-1 on *in silico* docking scores and, most notably, achieved a 12% *in vitro* hit rate across 21 unique scaffolds versus 3% for BoltzMol-1. FlexBench thus provides a framework for assessing structure-based AI methods for SMDD.

## 2 Methods

FlexBench includes two recent structure-based AI methods for SMDD: (1) SAGE-Flex, a model we built specifically for SMDD, and (2) BoltzMol-1 [7], an adaptation of Boltz-2 [14] for SMDD. These methods were compared against virtual screening and two versions of AI-accelerated docking.

**Virtual Screening** is a standard computational drug discovery method that uses physics-based molecular docking to score a large library of molecules against a target protein [2]. Here, FlexBench used Glide SP [1] to dock a random sample of 700,000 Enamine REAL Space molecules [4], which were then filtered to the top 20,000 by docking score. See Appendix A.

**Flex Active Learning** is an AI-accelerated docking method that uses the open-source graph neural network Chemprop [33–35] in an active learning loop [5] to accelerate Glide SP docking [1]. Over five iterations, it alternates between training Chemprop on Glide SP docking scores and applying the trained Chemprop model to select new Enamine REAL Space molecules for Glide SP docking. It then returns the top 20,000 molecules by Glide SP score across iterations. See Appendix B.

**Generative Glide** [6] is an AI-accelerated docking method from Schrödinger that similarly trains a fast AI model on molecules scored by Glide SP, but instead of using it to score random molecules from a large library, it trains the generative model REINVENT [36] to generate SMILES strings of molecules within the Enamine REAL Space that are predicted to have good docking scores. We generated 90,000 molecules and filtered them to the top 20,000 by Glide SP score. See Appendix C.

**BoltzMol-1.** Boltz [11, 14] is a co-folding model based on AlphaFold 3 [10] that was originally an open-source tool at MIT and is now maintained by Boltz PBC. On the Boltz platform, BoltzMol-1 [7] generates small molecule binders by pairing a generative model over the Enamine REAL Space with Boltz-2’s affinity module as the reward function. We generated 10,000 molecules per target, which BoltzMol-1 filtered internally to around 1,000 molecules using structural alerts. See Appendix D.

**SAGE-Flex** is a diffusion model [37] built internally and trained on the Protein Data Bank (PDB) [38–40] for SMDD. Like its predecessor MedSAGE [41], SAGE-Flex represents the ligand and protein residues as common chemical fragments to reduce the dimensionality of the complex and increase drug-likeness. Unlike MedSAGE, which assumes a rigid pocket, SAGE-Flex co-generates the small molecule and the pocket, thereby modeling an induced fit of the ligand to the pocket. The diffusion model generated 1–1.5 million compounds, which were filtered by Glide HTVS (a faster but less accurate version of Glide SP) to the top 5,000. These top molecules were then mapped to 2–3 million Enamine REAL Space analogs, which were subsequently filtered with Glide HTVS to select the top 20,000 molecules. SAGE-Flex thus generates compounds via a diffusion model and evaluates them with an orthogonal physics-based scoring function, combining the creativity of structure-based generative AI with the robustness of docking. See Appendix E.

## 3 *In Silico* Experiments

### 3.1 Setup

#### Targets

FlexBench includes four internal targets: three class A GPCRs (GPCR A1, A2, A3) with orthosteric pockets and one class B GPCR (GPCR B1) with an allosteric pocket. All methods had access to the same holo (small-molecule-bound) structure and Glide grid, although not all methods used them. GPCR A1, A3, and B1 used PDB structures while GPCR A2 used a homology model.

#### Compute Budget and Chemical Space

All methods used approximately $200 of compute per target on Amazon Web Services CPU and/or GPU instances or on the Boltz platform. Methods could generate any number of molecules internally but had to output at most 20,000 molecules to limit evaluation costs. Methods were restricted to the Enamine REAL Space [4] v2025.02 with 78.1 billion molecules.

#### Filtering

Because synthesis and *in vitro* testing are expensive, we filtered the molecules output by each method (“prefilter”) to the top 200 molecules (“postfilter”) using molecular weight, number of rotatable bonds, log solubility as predicted by ADMET-AI [42], Glide SP score, strain energy, blocked hydrogen bond donors, and diversity via Tanimoto clustering.

### 3.2 Results

Below, we describe results across all five methods and four targets focused on four areas of evaluation: chemical spaces, docking scores, diversity, and novelty. Additional results are in Appendix F.

#### Chemical Spaces

One of the most distinct *in silico* results was that the structure-based AI methods explored largely unique chemical spaces compared to each other and to docking-based methods (Figure 2a). This was generally consistent across all four targets both qualitatively (Figure 4) and quantitatively based on Tanimoto similarity (Figure 6). This illustrates that structure-based AI methods can design new molecular scaffolds that are not readily identified by traditional docking-based methods, even though all methods had access to the same chemical space.

**Figure 2:**
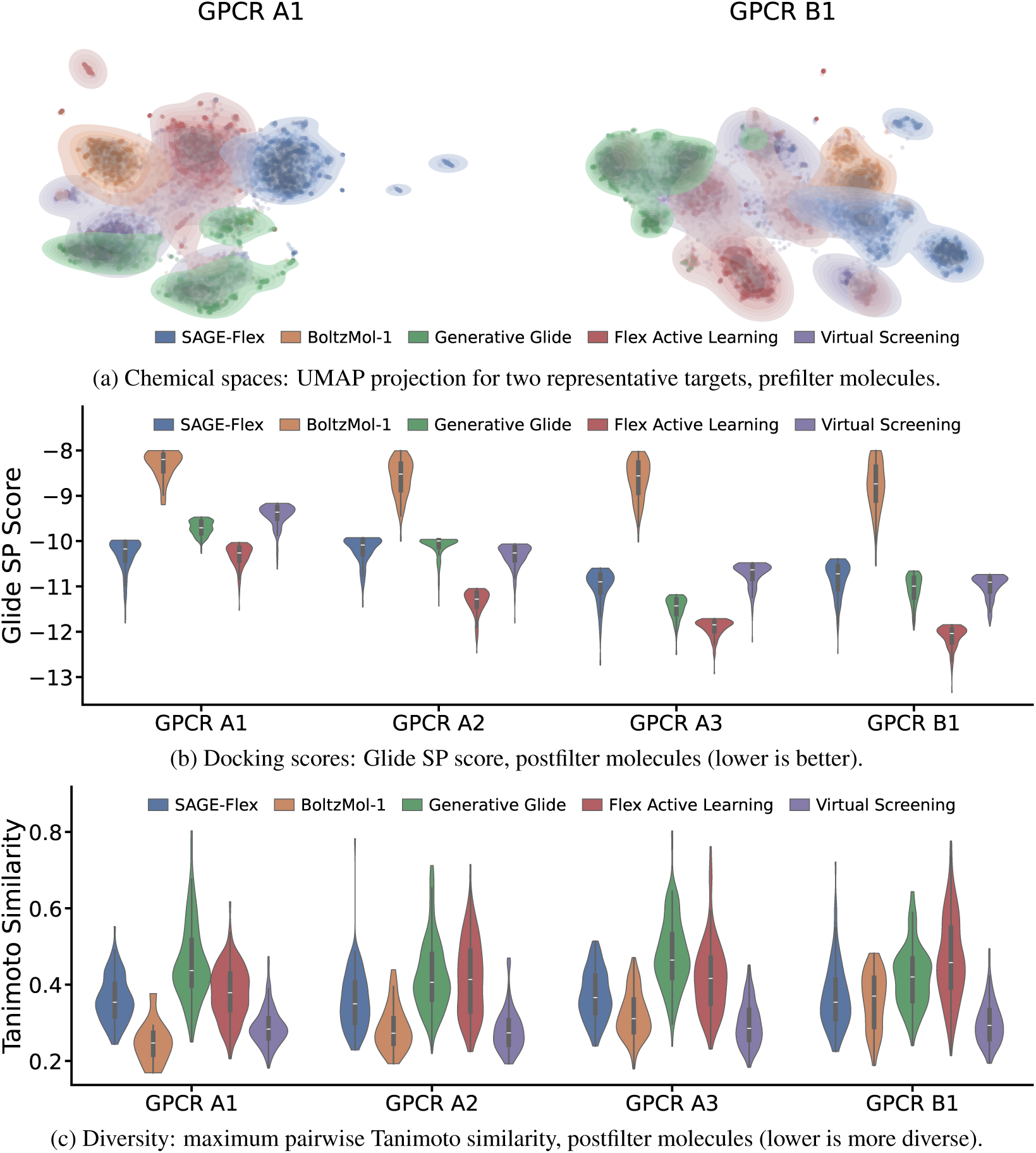
*In Silico* Results. Representative *in silico* metrics across methods and targets.

**Figure 3:**
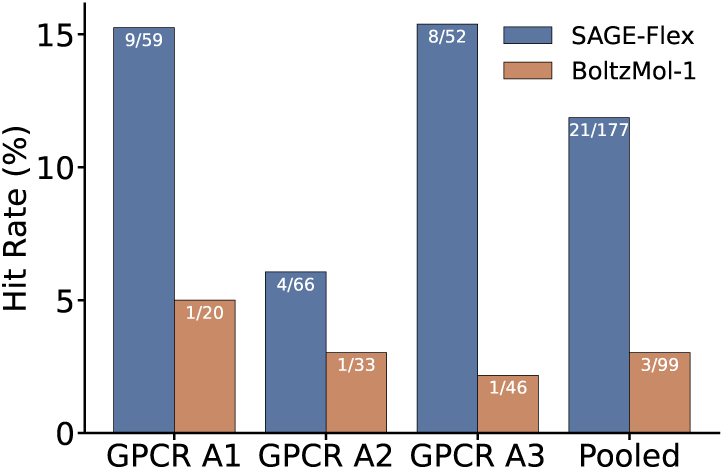
*In Vitro* Hit Rates. Hit rates for the top molecules from SAGE-Flex and BoltzMol-1 on three protein targets, and the pooled hit rate across the three targets.

**Figure 4:**
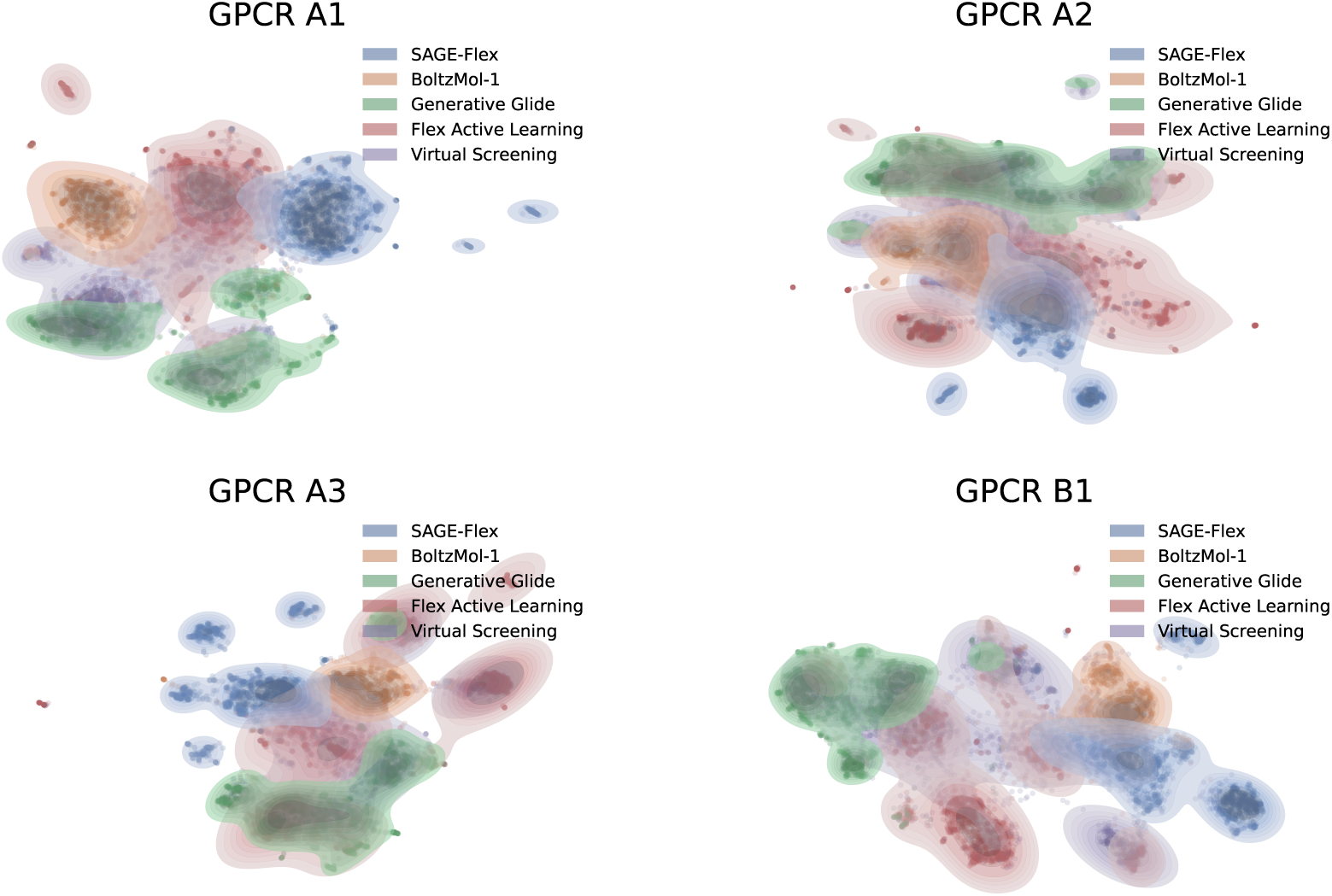
Chemical Spaces Across Targets, Prefilter Molecules. UMAP projection of each target’s chemical space, with a filled density per method. GPCR A1 and GPCR B1 also appear in Figure 2.

**Figure 5:**
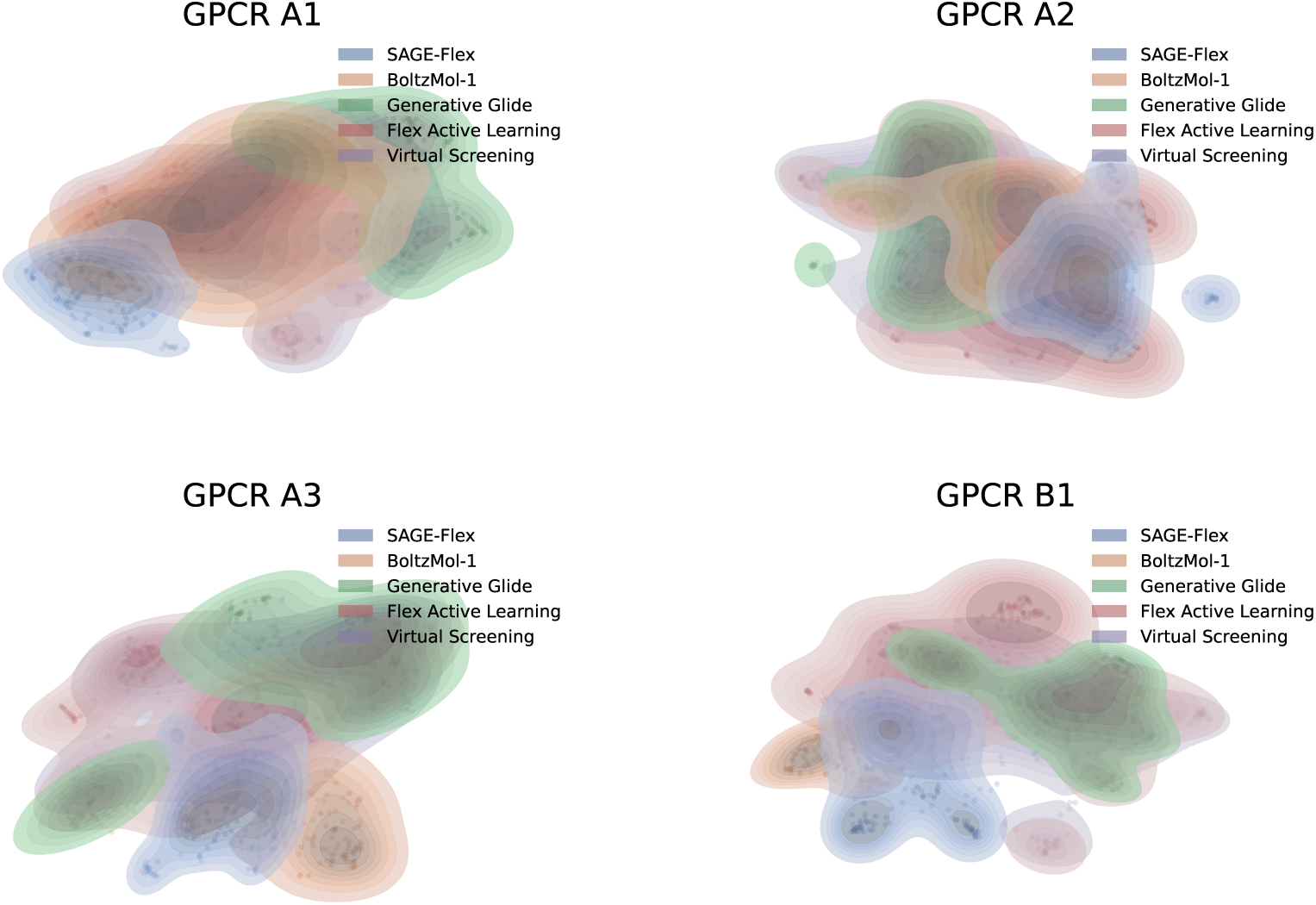
Chemical Spaces Across Targets, Postfilter Molecules. UMAP projection of each target’s chemical space, with a filled density per method.

**Figure 6:**
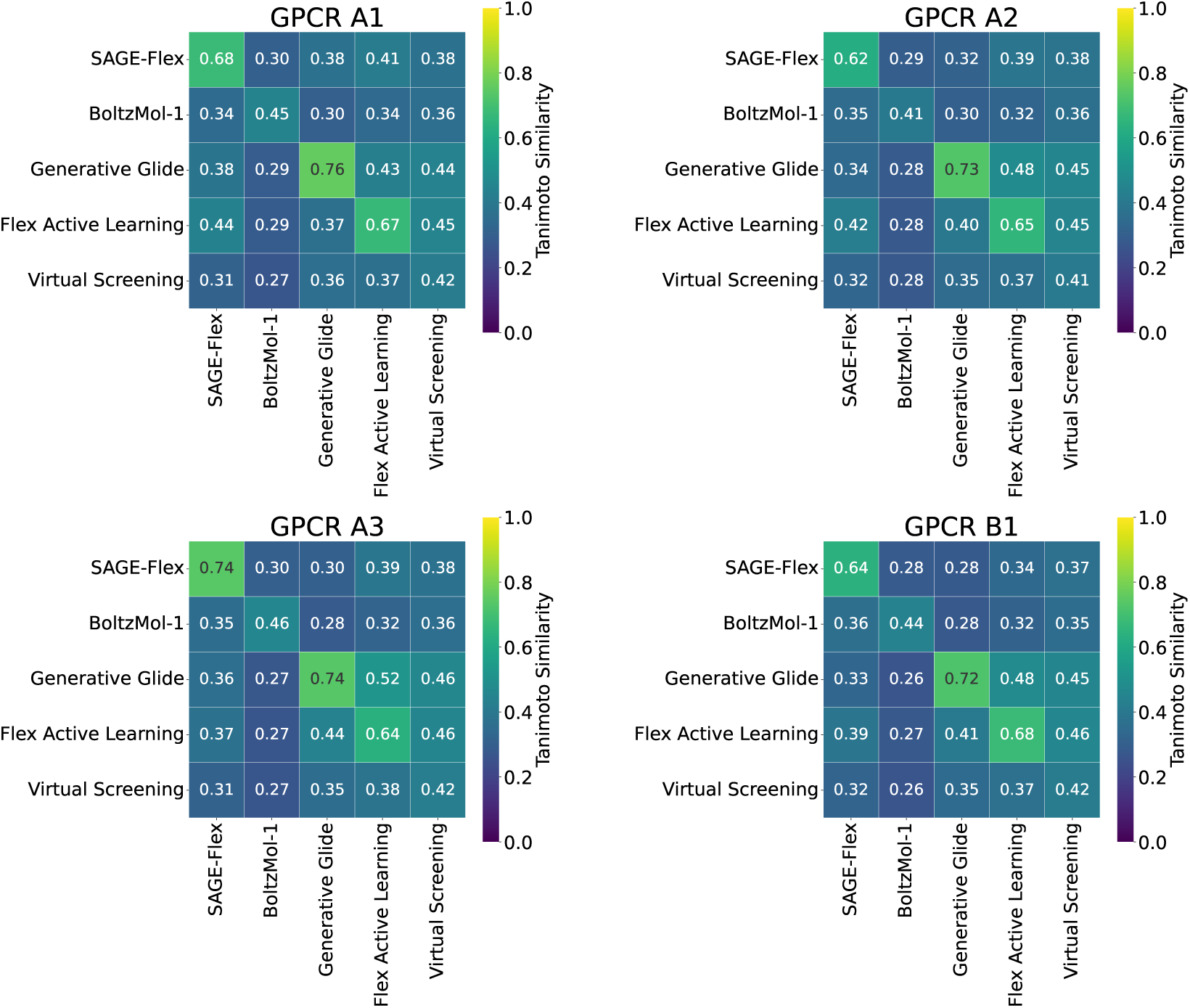
Between-Method Similarity Across Targets, Prefilter Molecules. For each pair of methods, the average over the row method’s molecules of each molecule’s maximum Tanimoto similarity to any molecule of the column method. The diagonal is the within-method value, excluding each molecule’s self-similarity.

#### Docking Scores

Designing new molecular scaffolds is only useful if they have a desired biological effect upon binding to the protein target, so we evaluated target binding potential by using Glide SP to generate poses and score molecules for each target. Because three methods optimized Glide SP directly and a fourth filtered on Glide HTVS, we also rescored the best Glide SP pose with AutoDock Vina [43], an independent scoring method. Under Glide SP (Figure 2b), all methods except for BoltzMol-1 generated molecules with good docking scores (all below −9 after filtering). Even when BoltzMol-1 was given a 5x larger compute budget (as recommended by Boltz), the BoltzMol-1 docking score distributions were unchanged despite up to 20x more molecules with a high Boltz confidence (Appendix G), indicating a gap between Boltz confidence and docking scores. Among the other methods, Flex Active Learning scored best on all four targets, while SAGE-Flex, Generative Glide, and Virtual Screening traded places in the middle. Under Vina (Figure 8), the methods were less differentiated, indicating overfitting to Glide SP, but the extremes were largely unchanged, with Flex Active Learning as the best and BoltzMol-1 as the worst on three of four targets each.

**Figure 7:**
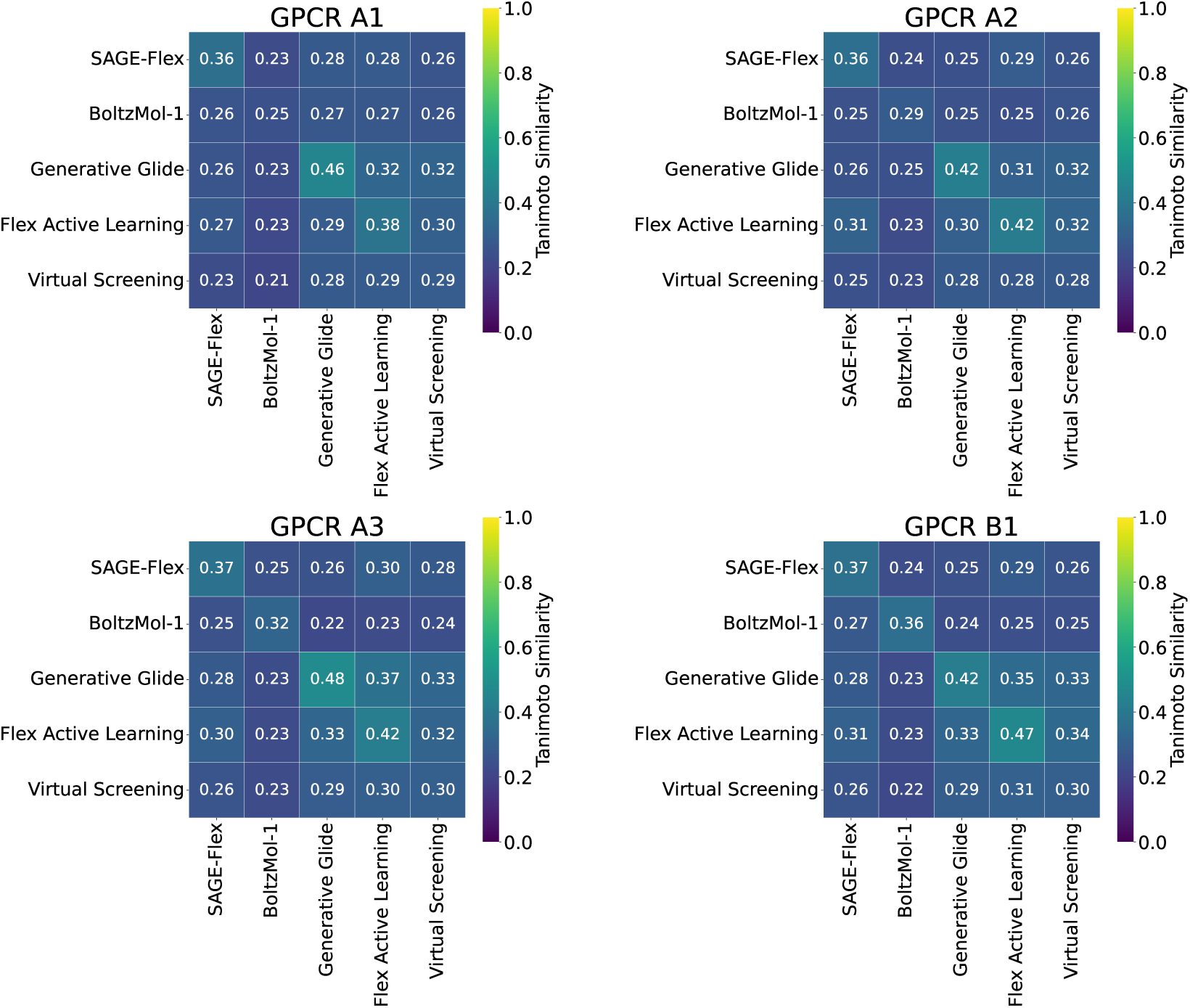
Between-Method Similarity Across Targets, Postfilter Molecules. For each pair of methods, the average over the row method’s molecules of each molecule’s maximum Tanimoto similarity to any molecule of the column method. The diagonal is the within-method value, excluding each molecule’s self-similarity.

**Figure 8:**
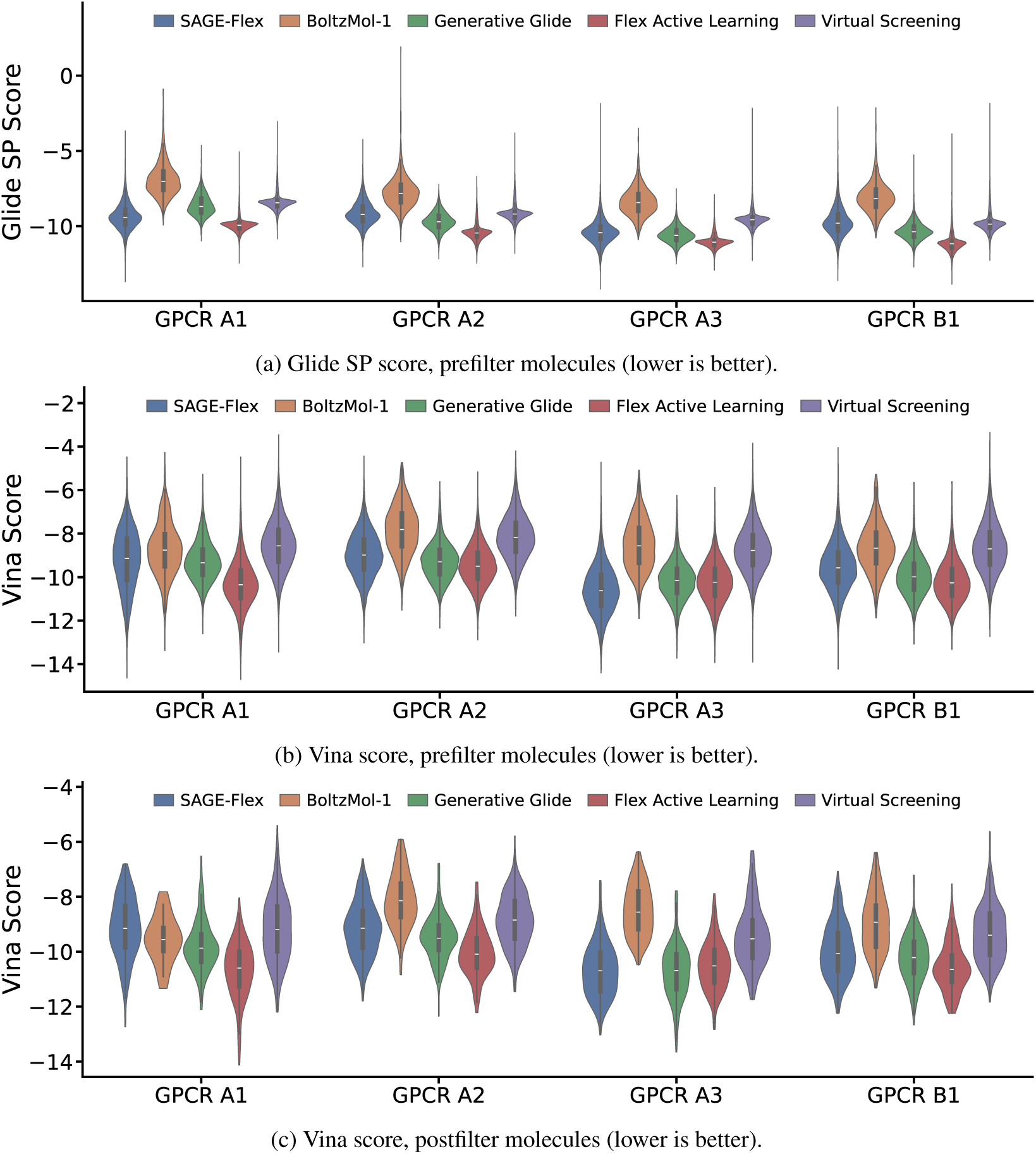
Docking Scores Across Targets. The docking score results that Figure 2 does not show: the prefilter Glide SP scores, and the Vina scores of the best Glide SP pose before and after filtering.

#### Diversity

Because docking scores are a weak proxy for binding, a diverse set of high-scoring molecules increases the chance of a hit and provides more options for lead optimization. We evaluated diversity by computing maximum pairwise Tanimoto similarities (Figure 2c) and number of unique scaffolds (Figure 9). BoltzMol-1 was an outlier since fewer than 200 of its molecules passed the filters, which inflated its apparent Tanimoto diversity. Among the others, Virtual Screening was the most diverse followed by SAGE-Flex and then Generative Glide and Flex Active Learning.

**Figure 9:**
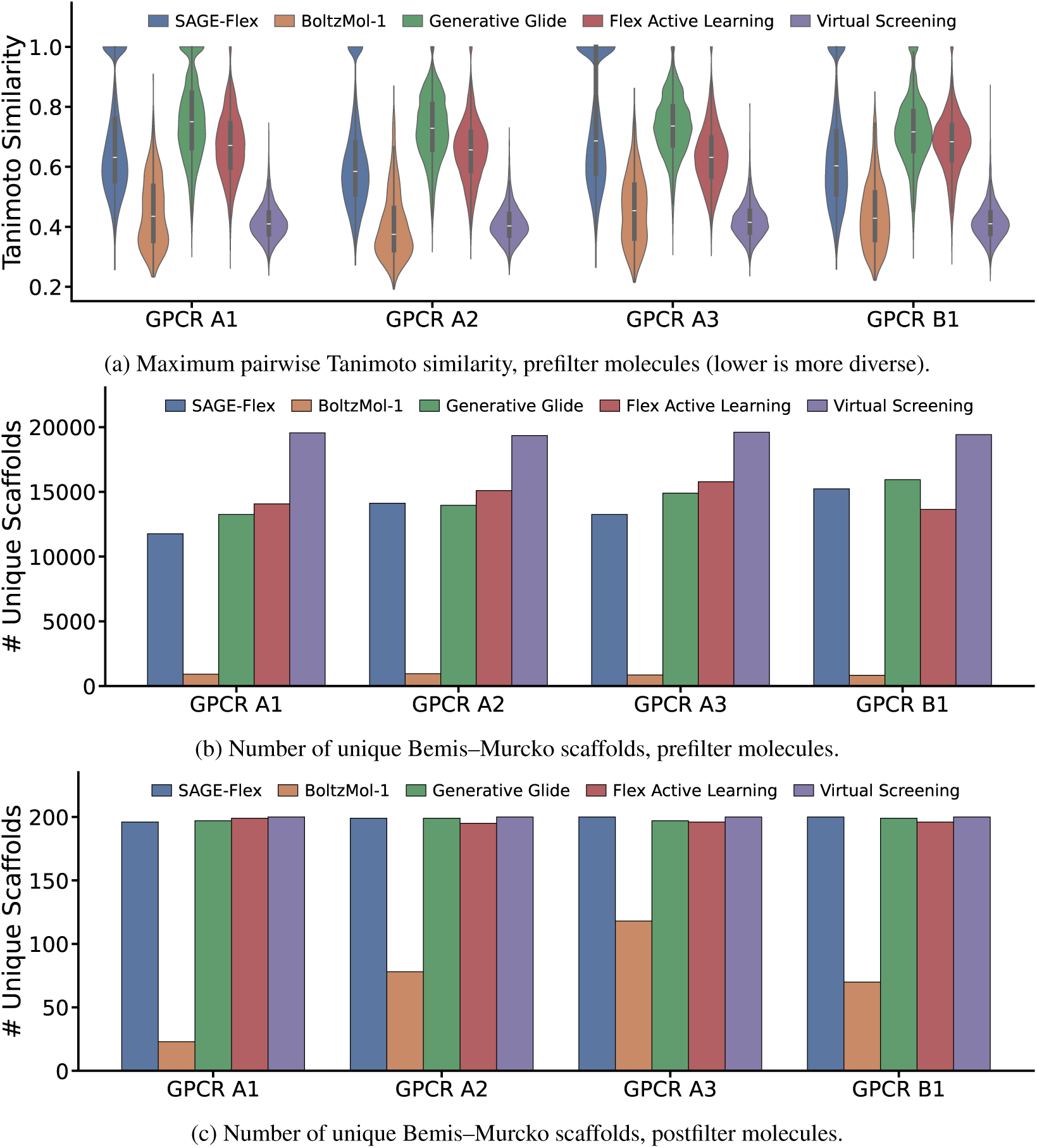
Diversity Across Targets. The diversity results that Figure 2 does not show: the prefilter maximum pairwise Tanimoto similarity, and the unique scaffold counts before and after filtering. Note that each method produced up to 20,000 molecules prefilter and 200 postfilter, except BoltzMol-1, which produced around 1,000 molecules, of which fewer than 200 molecules passed the filters.

#### Novelty

Since it is advantageous to have structurally novel chemical starting points (for proprietary intellectual property, to avoid known liabilities of previous scaffolds, etc.), we curated 3–54 patented molecules per target and computed each generated molecule’s Tanimoto similarity to them (Figure 10) and the percentage of novel scaffolds. On every target, all methods generated 100% novel scaffolds with low similarity to patented molecules, demonstrating sufficient novelty.

**Figure 10:**
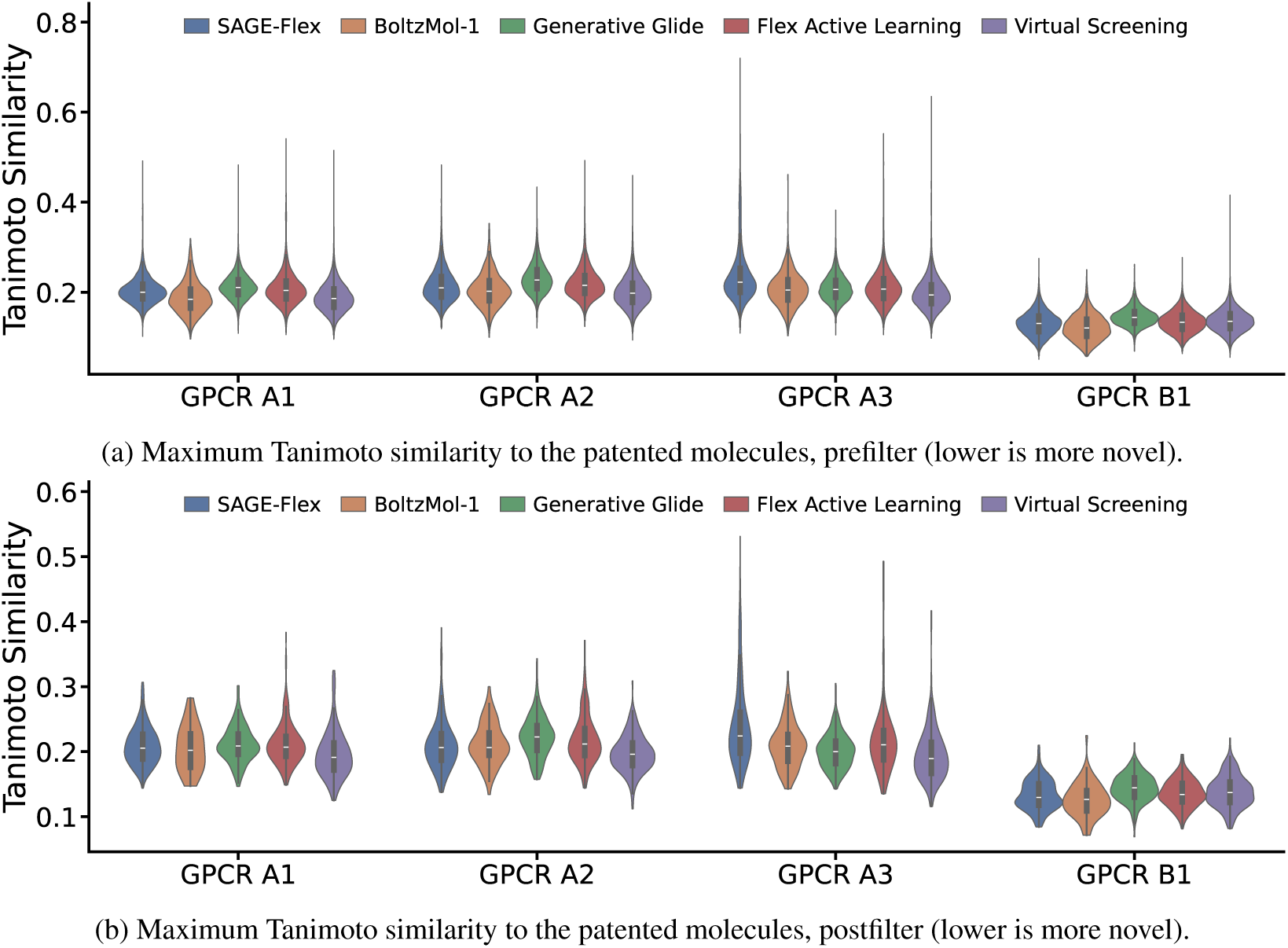
Novelty Across Targets. Each generated molecule’s maximum Tanimoto similarity to the patented molecules curated for its target, before and after filtering.

## 4 *In Vitro* Experiments

### 4.1 Setup

We compared the two structure-based AI methods, SAGE-Flex and BoltzMol-1, *in vitro* by testing 20–66 compounds per method per target (276 in total) depending on how many passed filtering and manual selection (Appendix H) and were successfully synthesized by Enamine (typically 80% within 3–4 weeks [44]). Docking-based methods have been previously validated *in vitro* [2, 3], so we did not test compounds from those methods due to cost and our focus on structure-based AI. The tested molecules came from earlier versions of SAGE-Flex and BoltzMol-1 with reduced compute budgets and a more limited filtering pipeline, so these results are conservative estimates of performance. Molecules were only tested *in vitro* on the class A GPCRs due to time and cost constraints.

GPCR A1 was assessed in a competition binding assay while GPCR A2 and A3 were assessed in functional assays run in antagonist mode (Appendix I). Hits were defined as molecules with *>* 50% inhibition at 10 *µ*M for GPCR A1 and A3. GPCR A2 proved harder with no molecules from either method exceeding 50% inhibition at 10 *µ*M, so we defined hits as molecules with *>* 50% inhibition at 50 *µ*M in the functional assay and *>* 25% inhibition at 50 *µ*M in an orthogonal binding assay.

### 4.2 Results

SAGE-Flex achieved a higher hit rate than BoltzMol-1 on all three targets (Figure 3), with a pooled hit rate of 12% compared to 3% (*p* = 0.01 pooling per-target odds ratios with the Cochran–Mantel–Haenszel test). Notably, the SAGE-Flex hits reached much higher percent inhibition than the single BoltzMol-1 hit in the GPCR A1 and GPCR A3 assays, while in the two GPCR A2 assays the BoltzMol-1 hit was comparable to the SAGE-Flex hits (Figure 17). The SAGE-Flex hits were also structurally diverse, with all 21 SAGE-Flex hits possessing different scaffolds.

## 5 Conclusion

We introduced FlexBench, a benchmark that compares, *in silico* and *in vitro*, structure-based AI and docking methods for SMDD. Each method achieved a different balance of *in silico* metrics, but the structure-based AI methods stood out by exploring regions of chemical space that were largely disjoint from the docking-based methods, enabling the design of new molecular scaffolds. SAGE-Flex in particular designed molecules with good docking scores in that new region of chemical space, demonstrating the value of a hybrid approach that combines structure-based generative AI to propose novel chemotypes with physics-based docking to evaluate them. This *in silico* success translated to the *in vitro* setting, where SAGE-Flex obtained a 12% hit rate across 21 distinct scaffolds versus 3% for BoltzMol-1. The current results are limited to four internal GPCR targets, and only two methods were tested *in vitro* in an earlier configuration. Moving forward, we plan to solidify the pipeline from *in silico* to *in vitro* results and extend the validation to more methods and public targets. By tying computational metrics to experimental outcomes under a fixed budget and chemical space, FlexBench provides a framework for evaluating structure-based AI models for small molecule drug design.

## Acknowledgments and Disclosure of Funding

Thank you to Jeremy Wohlwend, Geoffrey Smith, and Demitri Nava at Boltz PBC for their generous technical support and guidance as well as free credits on the Boltz platform. Thank you to Ben King, Nikita Chopra, Kun Yao, and Nancy Guan at Schrödinger for early access to Generative Glide as well as for technical insights and guidance for running Generative Glide and Glide SP. All funding is provided by Flex Therapeutics.

## A Virtual Screening Additional Details

We ran virtual screening using a typical pipeline with tools from Schrödinger Release 2026-2^1^ as follows:

1. We built a Schrödinger grid file that defines the protein’s pocket.
2. We randomly sampled a list of 700,000 SMILES from the Enamine REAL Database 10.4M set^2^, which is a diverse subset of the Enamine REAL Space.
3. We ran Schrödinger’s LigPrep on the SMILES to prepare them for docking. This included expanding tautomeric and ionization states, ring conformations, and stereoisomers.
4. We ran Schrödinger’s Glide SP [1] using the protein grid file and the ligands prepared by LigPrep.
5. We ran Schrödinger’s Glide Merge across the outputs of Glide SP to keep only the top 20,000 ligands by docking score.

## B Flex Active Learning Additional Details

Flex Active Learning is an active learning approach that takes advantage of the relative accuracy of Glide SP and the relative speed of the graph neural network (GNN) model Chemprop. Glide SP takes roughly 10 seconds per molecule [45], while Chemprop is about 5 orders of magnitude faster, taking around 0.00015 seconds per molecule (based on internal experiments using Chemprop on an AWS gr6.4xlarge instance with one NVIDIA L4 GPU and 16 vCPUs). Chemprop can therefore search far more of chemical space than Glide SP alone, with Glide SP reserved for determining the true docking scores of the most promising molecules that Chemprop identifies. On each iteration, Chemprop gets more accurate as it trains on more data points with better scores from Glide SP.

Flex Active Learning worked as follows. First, we randomly sampled 25,000 molecules from the Enamine REAL Database 10.4M set, and we scored them with Glide SP. Then, we ran five active learning iterations:

1. **Train:** We trained Chemprop on all molecules scored with Glide SP so far.
2. **Predict:** We used Chemprop to make predictions on 500M molecules randomly sampled from the Enamine REAL Database 10.4B set. Note that this is the 10.4B set, not the 10.4M set used for the initial sample, so it contains 1000x more molecules. Also note that we used a distinct set of 500M molecules on each round.
3. **Filter:** We filtered the 500M molecules to the top 25,000 molecules by predicted docking score.
4. **Evaluate:** We ran Glide SP on the top 25,000 molecules to determine true docking scores.

Finally, after completing these five iterations, Flex Active Learning returned the top 20,000 molecules by docking score from the 150,000 molecules scored by Glide SP.

## C Generative Glide Additional Details

For Generative Glide, we used the default settings except that we trained the docking score prediction AI model on 50,000 data points for 30 hours instead of 20,000 data points for 12 hours, we used 32 generative agents instead of 20, and we generated 90,000 molecules instead of 20,000. These molecules were filtered to the top 20,000 molecules based on Glide SP docking scores.

## D BoltzMol-1 Additional Details

We applied BoltzMol-1 using the Boltz platform (https://lab.boltz.bio) to generate 10,000 small molecules per target. At the suggestion of scientists at Boltz PBC, we also experimented with generating 5x as many Boltz molecules (i.e., 50,000 molecules instead of 10,000). Because the model learns during generation, this experiment was expected to generate results that would be more than 5x better. Since it exceeded our compute budget, we defer its results to Appendix G. In both cases, BoltzMol-1 then performed internal filtering using constraints such as structural alerts, resulting in a final output of around 1,000 molecules per target.

### Structure and pocket definition

BoltzMol-1 folds its own structure from the protein sequence rather than taking PDB coordinates as input, so it does not consume the shared holo structure in the same form as the other methods. It was nevertheless given the same structural information. On the Boltz platform, we specified the binding site for each target using the pocket residues and an example bound ligand read directly off the same structure that every other method received, and we then confirmed for each target that the resulting Boltz-predicted structure agreed with that holo structure and that the predicted pocket was the intended one. The exception was GPCR B1, the class B allosteric target, where sequence-based prediction alone did not reliably place the pocket; there we additionally supplied the holo PDB structure as a template. In all cases, therefore, BoltzMol-1’s structure and pocket definition derived from the same structure used by the other methods, even though the coordinates it designed against were its own prediction rather than the deposited ones.

## E SAGE-Flex Additional Details

SAGE-Flex generated molecules through a multi-stage workflow that combined fragment-based generative modeling, docking-based prioritization, and mapping into synthetically accessible chemical space. First, SAGE-Flex used a diffusion model to generate approximately 1–1.5 million compounds directly within the target binding site. Rather than generating molecules atom-by-atom, SAGE-Flex operated at the level of molecular fragments, iteratively selecting and positioning ligand fragments within the protein pocket, while adapting the pocket conformation simultaneously. Fragments were represented using learned embeddings that encode both their chemical structure and molecular properties, as described in the MedSAGE framework [41]. Both the protein and the growing ligand were represented using this latent fragment-level representation.

The generated compounds were then docked using Glide HTVS, and the top 5,000 molecules were retained based on docking score. Although these generated molecules were designed to be drug-like and chemically reasonable, they were not necessarily directly purchasable within the Enamine REAL Space. We therefore applied an internally developed method, Remix, to map the generated designs into accessible chemical space. Remix first fragmented each generated molecule, identified structurally and chemically similar building blocks available within the Enamine REAL collection, and then recombined selected building blocks using a fixed library of supported synthetic reactions.

Applying Remix to the 5,000 prioritized SAGE-Flex molecules typically produced 2–3 million candidate analogs in the Enamine REAL Space. These analogs were subsequently evaluated with Glide HTVS, and the highest-ranking 20,000 compounds were selected for downstream analysis and visual inspection. Importantly, the full computational workflow—including SAGE-Flex diffusion-based generation, Remix enumeration, and both rounds of Glide HTVS docking—was included within the approximately $200 compute budget per target.

## F Additional *In Silico* Results

Here we present all *in silico* results that are not included in Figure 2 of the main text.

### F.1 Chemical Spaces

Figures 4 and 5 show the UMAP projection of every target’s chemical space, and Figures 6 and 7 the average maximum Tanimoto similarity between each pair of methods.

Prefilter, the pattern described in Section 3 was visible on all four targets: the structure-based AI methods occupied regions the docking-based methods did not reach, the AI-accelerated docking methods overlapped with Virtual Screening, and every method’s within-method similarity exceeded all of its between-method values.

Postfilter, each method was down to at most 200 molecules, which was too few for a density estimate to resolve the same structure. The between-method similarities did still separate the methods, with each one’s within-method value generally exceeding its between-method values.

### F.2 Docking Scores

Figure 8 shows the prefilter Glide SP scores and the Vina scores before and after filtering. Filtering shifted every method’s Glide SP distribution lower, since a target-dependent Glide SP threshold was one of the filters, but it did not reorder which method was best or worst: Flex Active Learning scored best and BoltzMol-1 worst on all four targets, prefilter and postfilter alike. Under Vina the methods compressed toward each other, and the two extremes held on three of four targets each rather than on all four, as discussed in Section 3.

### F.3 Diversity

Figure 9 shows the prefilter maximum pairwise Tanimoto similarity and the unique scaffold counts before and after filtering. Prefilter, nearly all Virtual Screening molecules had different scaffolds, while SAGE-Flex, Generative Glide, and Flex Active Learning had fewer and had more similar molecules. Postfilter, every method had essentially one scaffold per molecule, but they still differed in diversity according to the pairwise similarities (Figure 2). Filtering lowered every method’s pairwise similarity, since Tanimoto clustering was one of the filters, and it generally preserved the ordering seen prefilter. As a reminder, BoltzMol-1 was an outlier for similarity calculations since it generated around 1,000 molecules instead of 20,000 and fewer than 200 of its molecules passed all the filters, which inflated its apparent diversity.

### F.4 Novelty

Figure 10 shows novelty with respect to the patented molecules curated for each target. Both prefilter and postfilter, the generated molecules were novel across all methods and targets with 100% unique scaffolds compared to the patented molecules and generally low Tanimoto similarity to them.

### F.5 Molecular Properties

Figures 11 and 12 show molecular weight, logP, QED drug-likeness, and the percentage of molecules free of PAINS alerts, before and after filtering.

**Figure 11:**
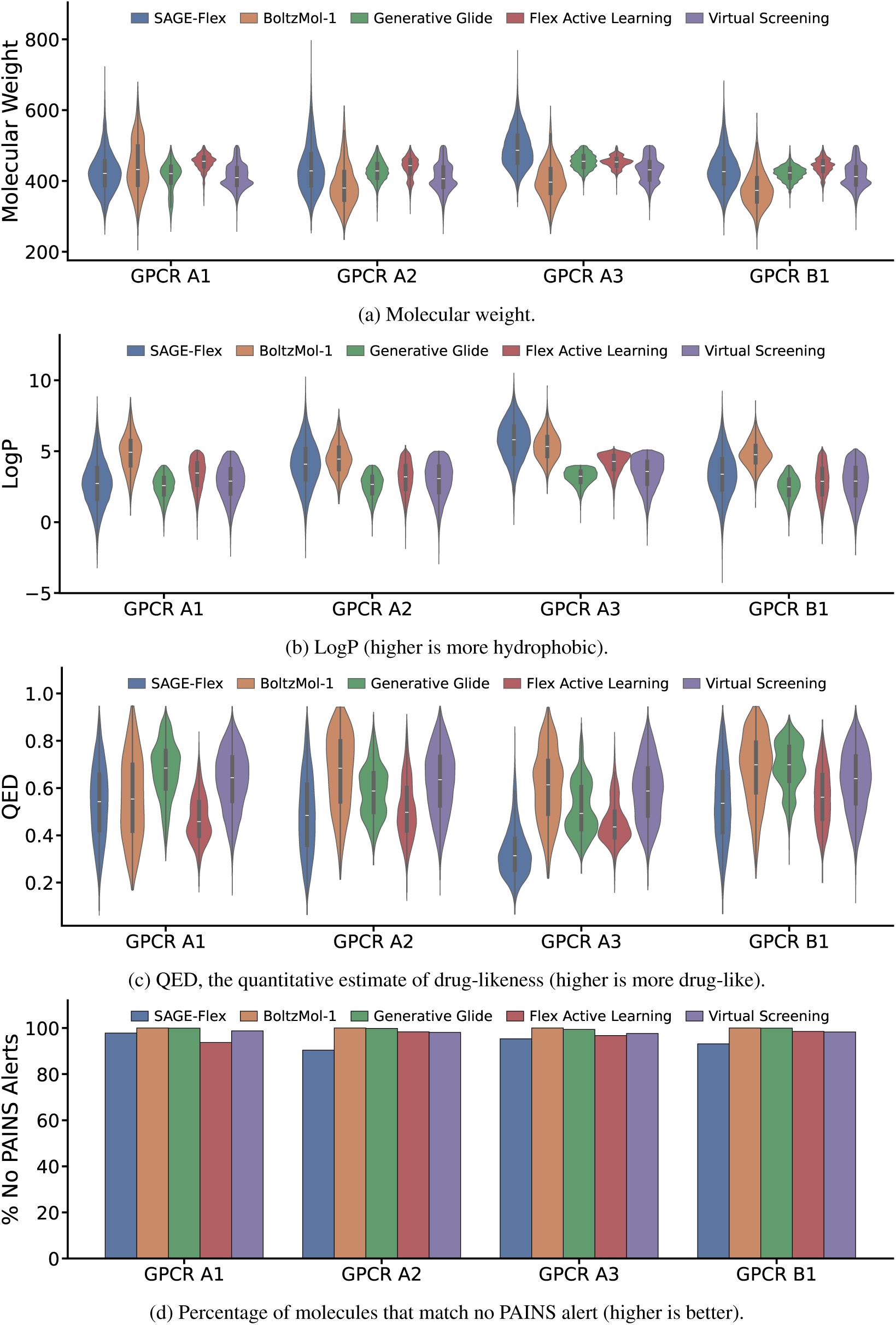
Molecular Properties Across Targets, Prefilter Molecules. Molecular weight, logP, QED drug-likeness, and freedom from pan-assay interference compound (PAINS) alerts, for the molecules generated by each method.

**Figure 12:**
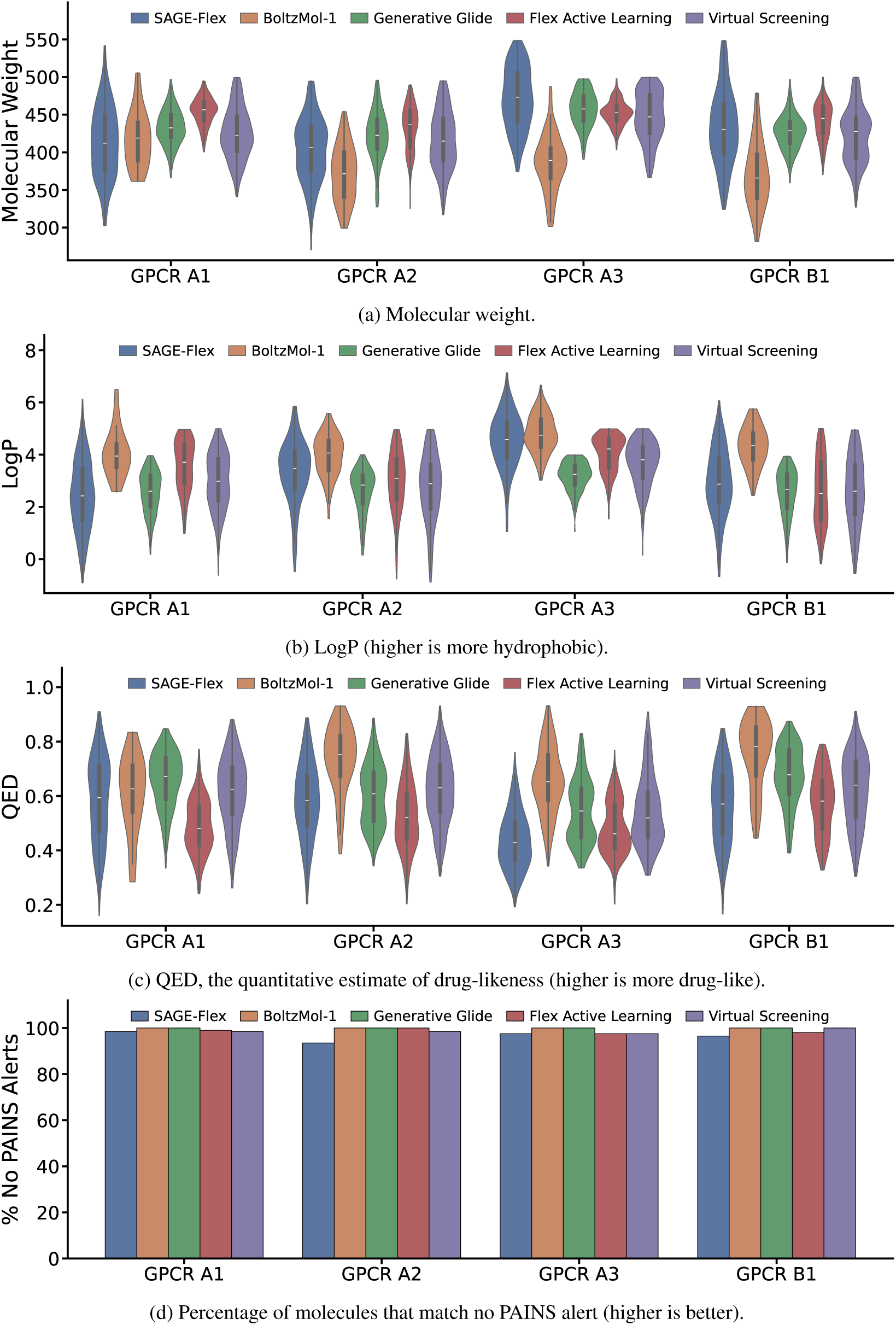
Molecular Properties Across Targets, Postfilter Molecules. Molecular weight, logP, QED drug-likeness, and freedom from pan-assay interference compound (PAINS) alerts, for the molecules generated by each method.

## G BoltzMol-1 5x Experiment

In our main experiments, we ran BoltzMol-1 to generate 10,000 molecules for each target in order to satisfy our fair compute budget constraint. However, since BoltzMol-1 learns as it generates molecules, the scientists at Boltz PBC suggested that we run BoltzMol-1 for 5 times as many steps (50,000 molecules) to see if it would improve performance. We ran this experiment for the three class A GPCR targets. For the class B GPCR target, supplying the holo PDB structure as a template was enough to place the predicted pocket correctly in general (Appendix D), but BoltzMol-1 with 10,000 molecules still struggled to place the generated molecules within that pocket in a reasonable conformation, so we did not run the 5x experiment for that target.

According to Figure 13, which shows the trajectory of Boltz binding confidence over time for the generated molecules, running BoltzMol-1 for five times longer yielded more than five times as many high-confidence molecules. For example, for GPCR A1, the number of molecules with a binding confidence *>*0.6 increased from *∼*100 to *∼*2,000, representing a 20-fold increase for just 5 times more compute. The result was similar for GPCR A2, although for GPCR A3 the increase was less than 10-fold, perhaps because the model already generated a large number of high-confidence molecules in the first 10,000 steps.

**Figure 13:**
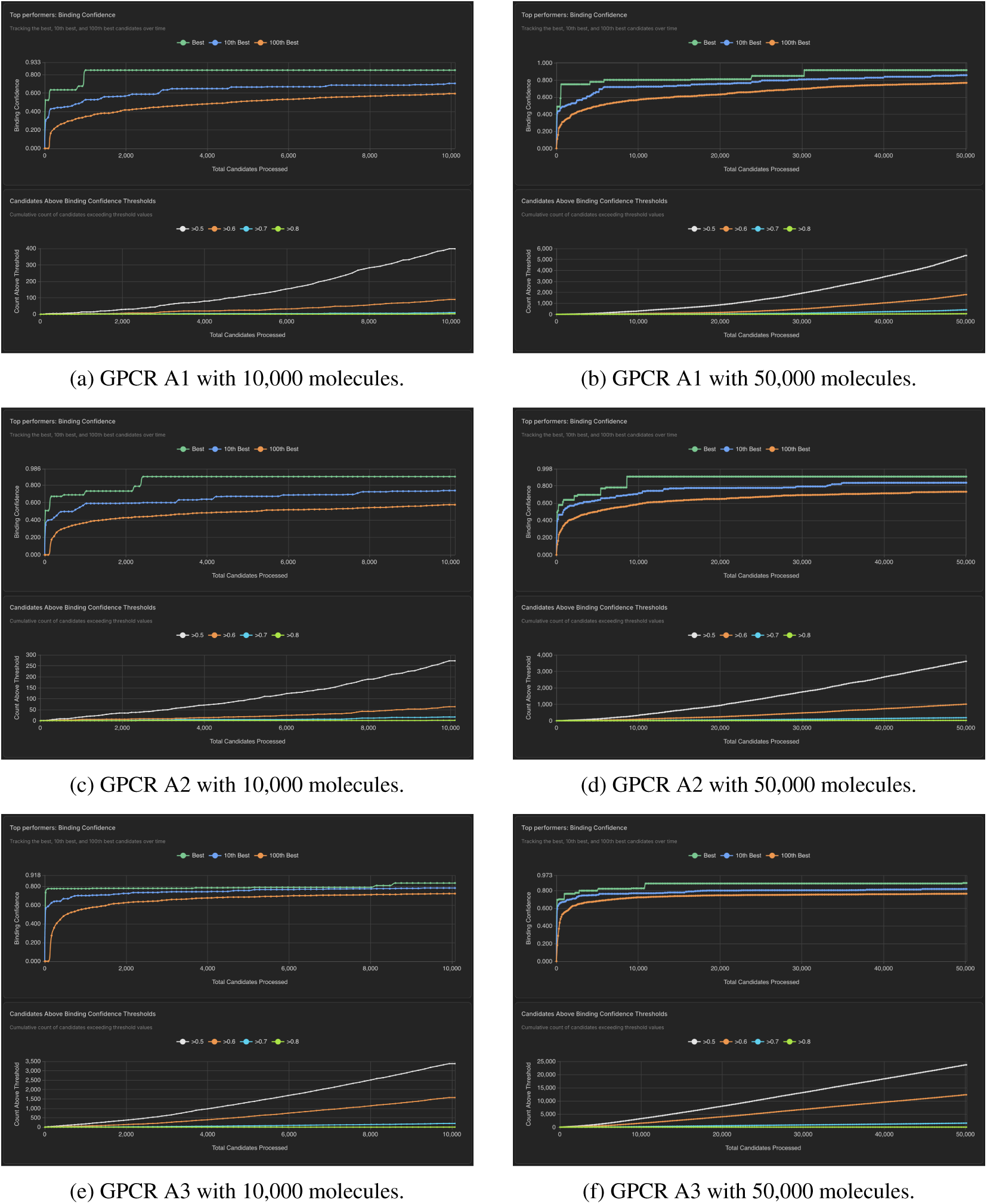
BoltzMol-1 5x Binding Confidence. Binding confidence of the molecules generated by BoltzMol-1 as a function of the number of candidates processed, comparing 10,000 (left) and 50,000 (right) generated molecules for each of the three class A GPCR targets. Each panel tracks the binding confidence of the best, 10th-best, and 100th-best candidates over time (top) and the cumulative number of candidates exceeding binding confidence thresholds (bottom).

However, the dramatic improvement in Boltz binding confidence values did not translate to improved docking scores. As shown in Figure 14, the distributions of docking scores (both Glide SP and Vina) for the 10,000- and 50,000-molecule runs were very similar. Furthermore, as seen in Figure 15, the diversity of the molecules from the 50,000-molecule run decreased, presumably because it focused on molecules with high Boltz binding confidence. The analysis of the chemical spaces of the two runs in Figure 16 shows that the two runs were similar, but the 50,000-molecule run did explore areas of chemical space beyond that of the 10,000-molecule run.

**Figure 14:**
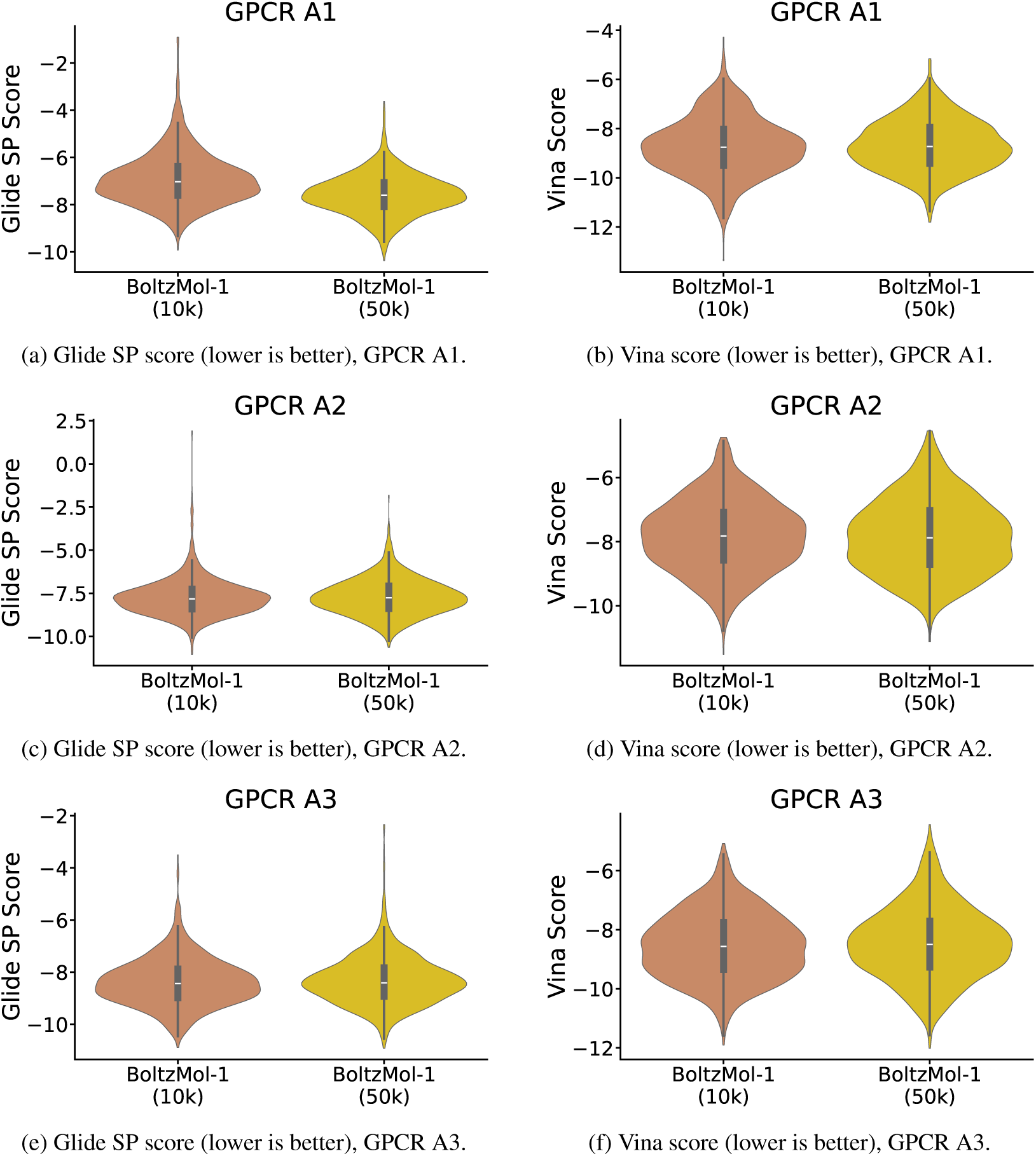
BoltzMol-1 5x Docking Scores. Distributions of prefilter docking scores for BoltzMol-1 with 10,000 and 50,000 generated molecules across the three class A GPCR targets.

**Figure 15:**
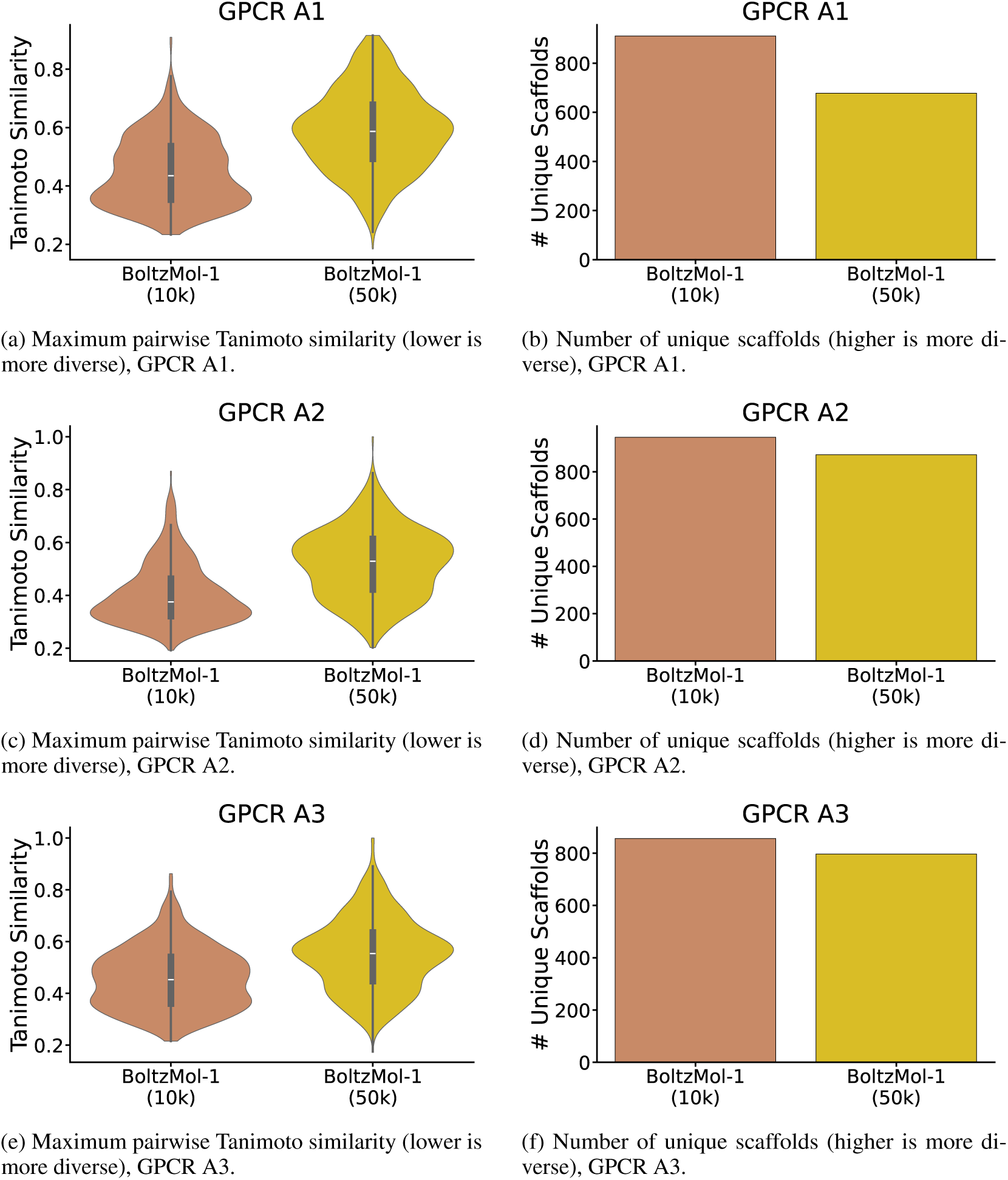
BoltzMol-1 5x Diversity. Diversity of prefilter molecules generated by BoltzMol-1 with 10,000 and 50,000 generated molecules across the three class A GPCR targets.

**Figure 16:**
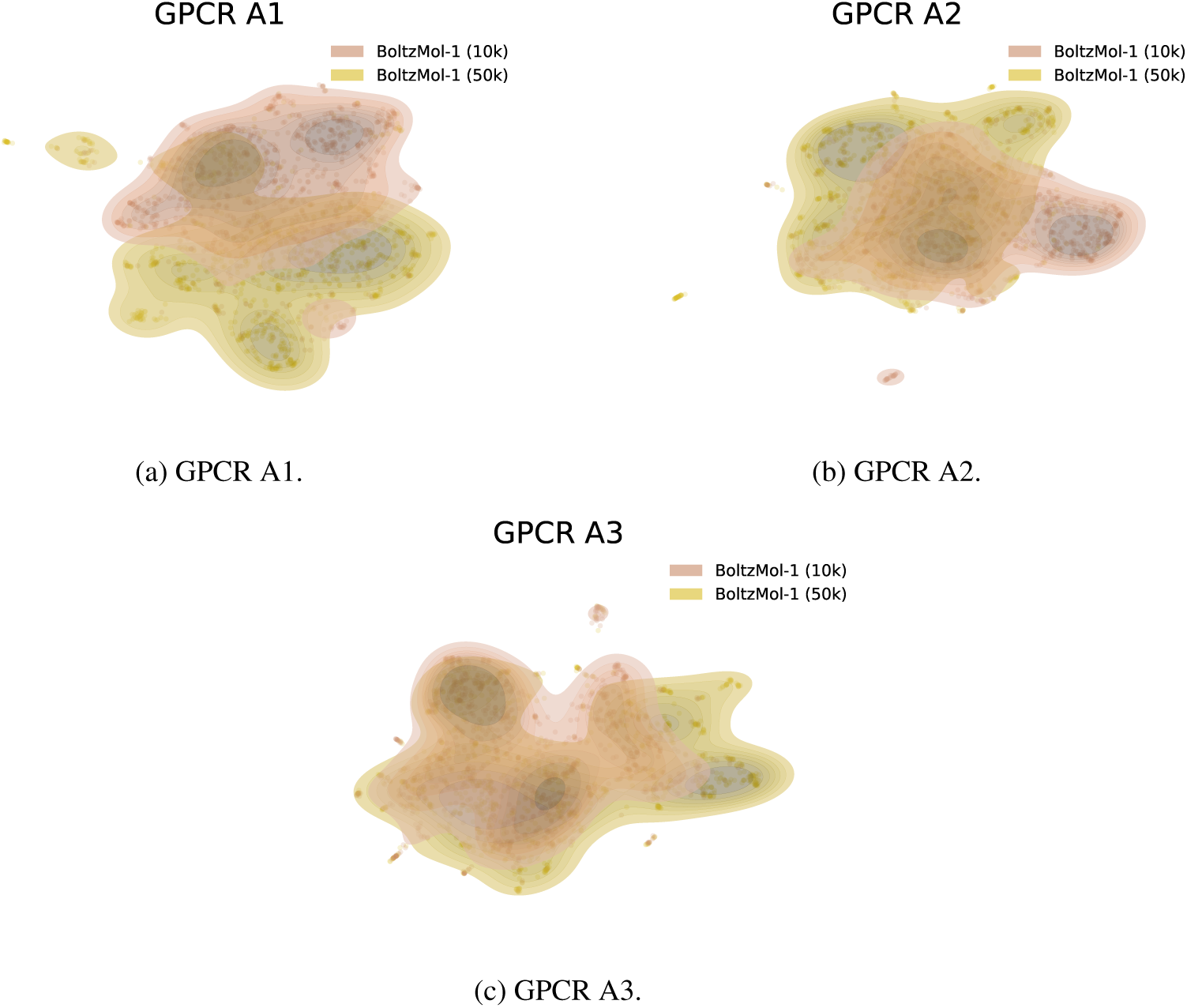
BoltzMol-1 5x Chemical Space. UMAP projections of the chemical space explored by prefilter molecules generated by BoltzMol-1 with 10,000 and 50,000 generated molecules across the three class A GPCR targets.

**Figure 17:**
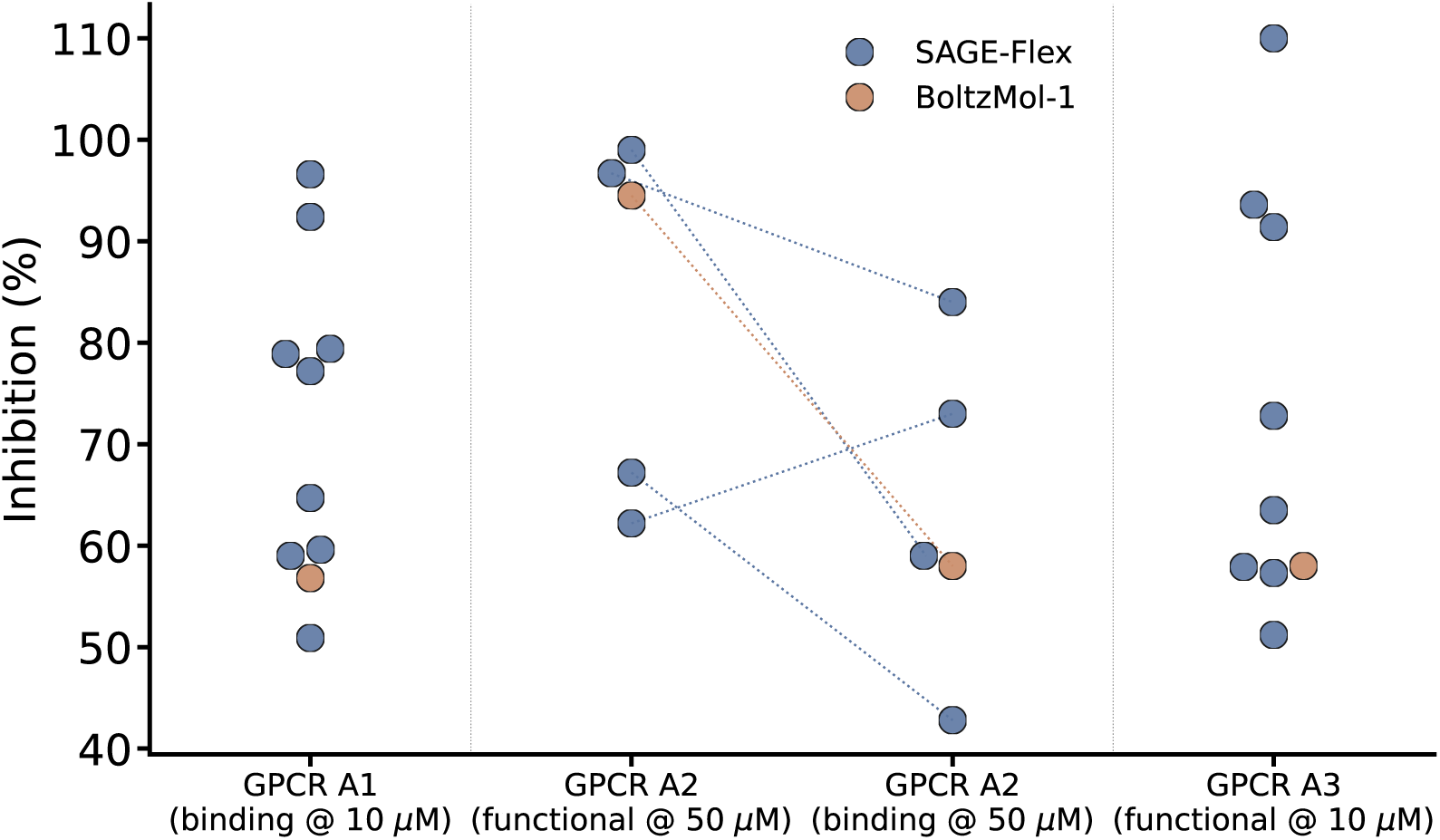
*In Vitro* Hit Activity. Percent inhibition of each individual hit from SAGE-Flex and BoltzMol-1, with one point per hit. GPCR A1 and GPCR A3 each have a single assay, while GPCR A2 hits appear in two sections—the primary functional assay and the secondary orthogonal binding counter-screen—with dotted lines connecting the two measurements of the same molecule. Assay details are in Appendix I.

Taken together, the outcome of the 5x experiment suggests that if the Boltz binding confidence is accurate, then the 50,000-molecule run is preferable due to its concentration on compounds with high binding confidence. However, the relatively poor docking scores and decreased diversity caution that as BoltzMol-1 generates more molecules, it may be concentrating on a small segment of chemical space that may not be optimal for the target.

## H Manual Selection

When finalizing the list of compounds for synthesis, we visually inspected the docked pose of each candidate that survived the automated filtering pipeline (Section 3) and deprioritized compounds that exhibited one or more of the following features:

- Errors introduced during 3D building or ligand preparation, including implausible tautomeric or protonation states, particularly where p*K*_a_ prediction is uncertain.
- Poor pocket complementarity, including “floating” poses that leave large gaps or unoccupied regions of the pocket and may bury energetically unfavorable or frustrated waters.
- Unsatisfied polar or charged groups, including hydrogen bond donors that are buried in hydrophobic parts of the pocket, in cases where this is not captured by the automated blocked hydrogen bond donor filter.
- Charged functional groups positioned in hydrophobic regions of the binding site. These criteria were applied identically to the molecules from SAGE-Flex and BoltzMol-1.

## I *In Vitro* Assay Protocols

Below we summarize the assays used for the three targets tested *in vitro*. GPCR A1 was assessed with a competition *binding* assay, while GPCR A2 and A3 were assessed with *functional* assays (in antagonist mode), which require a compound to modulate receptor signaling in cells and are therefore a more stringent readout than binding alone. For GPCR A2, an orthogonal radioligand *binding* counter-screen was run alongside the functional assay, and hits were required to be active in both. The cell-based assays were run in 384-well format using cell lines that overexpress the target receptor, and the membrane-based counter-screen in 96-well format. All compounds were tested in replicate wells.

### GPCR A1 (*binding*)

Cells expressing a terbium-labeled version of the receptor were incubated with a red fluorescent agonist tracer at a fixed concentration together with test compound at 10 *µ*M in triplicate for one hour at room temperature. The plate was read on a PerkinElmer EnVision using a homogeneous time-resolved fluorescence (HTRF) module. Compounds were assessed by their ability to compete with the tracer and reduce the HTRF signal, and activity is reported as percent inhibition of specific tracer binding.

### GPCR A2 (*functional*)

Calcium mobilization was measured on a FLIPR Tetra using the recombinant HEK293 stable pool expressing the receptor. Cells were plated at 12,000 cells/well the day before the assay and loaded with a calcium-sensitive dye for two hours at 37°C. The assay was run in antagonist mode: test compounds were pre-incubated with the cells at 10 *µ*M or 50 *µ*M in triplicate for 30 minutes at 25°C before challenge with a reference agonist at approximately its EC_80_ concentration. Activity is reported as percent inhibition of the agonist-stimulated calcium response.

### GPCR A2 (*binding counter-screen*)

Compounds were also profiled in an orthogonal radioligand competition binding assay. Membranes overexpressing the receptor (PerkinElmer) were incubated in duplicate with test compound at 50 *µ*M and a tritiated reference antagonist for the receptor in a Tris buffer containing MgCl_2_ and bovine serum albumin for two hours at 27°C. Bound radioligand was then captured on glass fiber filter plates, washed, and quantified by scintillation counting on a Revvity MicroBeta2 microplate counter. Activity is reported as percent inhibition of specific radioligand binding.

### GPCR A3 (*functional*)

cAMP accumulation was measured in antagonist mode using a HEK293 stable line overexpressing the receptor and a Cisbio HTRF cAMP detection kit (Cat# 62AM4PEJ). Test compounds were acoustically dispensed (100 nL) into 384-well plates at 10 *µ*M, to which *∼*1,000 cells per well were added in HBSS together with a fixed concentration of the reference agonist. Plates were incubated for one hour at 25°C. Intracellular cAMP was then quantified by adding cAMP-d2 and anti-cAMP detection reagents, incubating for a further hour at 25°C, and reading the HTRF ratio at 665/615 nm on a PerkinElmer EnVision; a cAMP standard curve (800 nM, 4-fold, 10 points) run on the same plate was used to convert HTRF ratios to cAMP concentrations. Activity is reported as percent inhibition of the agonist-stimulated cAMP response.

## J *In Vitro* Hit Activity

Figure 17 reports the measured percent inhibition of every *in vitro* hit; BoltzMol-1 contributed a single hit on each target while SAGE-Flex contributed several. In both GPCR A2 assays—the primary functional assay and the orthogonal binding counter-screen—the BoltzMol-1 hit was comparable in inhibition to the SAGE-Flex hits, but on the GPCR A1 binding assay and the GPCR A3 functional assay the SAGE-Flex hits reached substantially higher percent inhibition than the BoltzMol-1 hit.

## Footnotes

1 Schrödinger Release 2026-2: Glide, Schrödinger, LLC, New York, NY, 2026.

2 https://enamine.net/compound-collections/real-compounds/real-database-subsets

